# Micro and nanoplastic inhalation during pregnancy elicits uterine endothelial dysfunction in Sprague Dawley rats by impeding nitric oxide signaling

**DOI:** 10.1101/2025.09.12.675811

**Authors:** Chelsea M Cary, Taina L Moore, Andrew J Gow, Phoebe A Stapleton

## Abstract

Micro and nanoplastic (MNP) detection in human tissues demonstrates that exposure at any life stage is inevitable. We have previously demonstrated that pulmonary exposure to this emerging environmental contaminant impairs endothelial function in the uterine vasculature of nonpregnant and pregnant rats. However, neither the mechanism of this dysfunction nor the role of the endothelial-derived vasodilator, nitric oxide (NO), have been interrogated. Therefore, we assessed uterine macro- and microvascular reactivity in Sprague Dawley rats to determine the mechanistic role of NO signaling in endothelial dysfunction after repeated (gestational day 5-19) MNP inhalation during pregnancy. Results identified that MNP exposure reduced fetal growth and impaired endothelial-dependent dilation in the uterine microcirculation, which control placental perfusion and resource availability to the fetus. Levels of activated endothelial nitric oxide synthase (eNOS), phosphorylated on Ser^1176^, were substantially decreased (<50%) in uterine vessels from exposed rats. This suggests MNP inhalation limited NO production and bioavailability. Endothelial function was partially restored by supplementation of arterial segments with the eNOS cofactor tetrahydrobiopterin (BH_4_), demonstrating that exposed vessels were BH_4_-deficient. Partial restoration was also achieved by incubation with the reducing agent, DTT, suggesting that exposed vessels contained physiologically relevant levels of reactive oxygen and nitrogen species. Increased 3-nitrotyrosine residues and decreased thioredoxin protein expression further suggest MNP fosters nitrosative and oxidative stress in the uterine vasculature, impairing eNOS and endothelial-dependent dilation. These findings implicate eNOS uncoupling as a mechanistic basis for the vascular toxicity of MNPs and the adverse impact of MNPs on fetal development.

**NEW & NOTEWORTHY:** This study reveals that repeated micro and nanoplastic (MNP) inhalation throughout gestation blunts endothelial-dependent dilation in the uterine microcirculation, promoting fetal growth restriction. Exposure impaired endothelial nitric oxide signaling through deactivating endothelial nitric oxide synthase (eNOS), reducing the availability of the eNOS cofactor tetrahydrobiopterin and producing a nitrosative and oxidative environment in uterine vascular tissue. These novel findings highlight the eNOS uncoupling as a key mechanism behind the fetal growth restriction induced by MNP.

## INTRODUCTION

The environmental burden of micro and nanoplastics (MNPs) has risen rapidly over the last century. As plastic materials degrade, they break into tiny fragments that are within the micro (100nm< and >5mm) or nano (<100nm) size range. Within MNPs exists a fraction of small, lightweight particles (<10 µm) that become aerosolized and inhaled. MNPs have been identified in indoor and outdoor air (1-3), indoor and outdoor dust (4, 5), and the human lung (6, 7). Therefore, MNPs represent an inhalable component of particulate matter (PM). Epidemiological studies have linked PM inhalation to lethal cardiovascular impairments and events, including vascular dysfunction (8-10) and heart attack or stroke (11, 12). Controlled laboratory experiments in humans and animals have recapitulated these outcomes, reporting that particle inhalation disrupts vascular responses to stimuli in the macro- and microcirculation (13-17).

MNPs have been detected in the human placenta (18), which demonstrates MNP interaction with the uteroplacental vasculature. We have previously demonstrated that a single inhalation exposure to MNPs impairs vasodilation in the uterine vasculature of virgin female rats (19) and pregnant rats (20), which has negative implications for reproduction and development. Vascular responsivity, the ability of blood vessels to respond to stimuli, is paramount during pregnancy. Uteroplacental perfusion is tightly regulated to promote fetal growth and development. During pregnancy, the uterine arteries produce more signals to enable vasodilation and fewer signals that elicit vasoconstriction to deliver blood to the developing fetoplacental unit (21, 22). Impaired dilation of the uterine vasculature and reduced blood flow to the fetoplacental unit disrupts offspring development and cardiovascular physiology, as exemplified by reduced fetal weight, arterial stiffening, high blood pressure, and impaired vascular responsivity in experimental uterine artery ligation models (23-30). Similar health outcomes in offspring occur in models of maternal pulmonary particle exposure for which uterine arteriolar function is impacted (31-34).

Increased dilation of uterine arteries during pregnancy is achieved by elevated nitric oxide (NO) production, augmented prostacyclin release, and enhanced calcium signaling in the endothelium (35-39). This results in paracrine signaling to the vascular smooth muscle to stimulate relaxation, culminating in vasodilation, decreased vascular resistance, and increased blood flow. Elevated endothelial NO production is a key pregnancy-induced change in uterine vascular signaling with NO effects contributing to more than half of uterine artery blood flow (40). Similarly to uterine artery ligation models, inhibition of uterine NO signaling during gestation leads to decreased fetal weight (41-44), which suggests that vascular NO signaling is crucial for fetal development. Our previous work showed that a single pulmonary exposure to MNPs in late pregnancy caused fetal growth restriction and elicited endothelial-dependent dysfunction of the uterine radial artery, but not the major conduit vessel of the uterus, the uterine artery (20). The radial artery is a microvessel reliant on NO signaling that regulates placental perfusion to meet the metabolic demands of the placenta and fetus (45, 46). Its dysfunction, therefore, may cause relative ischemia in the fetoplacental unit. However, the role of NO-signaling changes in the uterine vasculature after MNP exposure during pregnancy remains unknown.

Pulmonary aerosol exposure is known to promote endothelial dysfunction and disrupted NO signaling through the scavenging of NO by radicals (47-49) and depletion of the cofactor, tetrahydrobiopterin (BH_4_) (50, 51) in the coronary, mesentery, skeletal muscle and dermal vasculature. These mechanisms, however, have only been marginally explored in the pregnant uterine vasculature. The effects of particle exposure and mechanisms at play are heavily dependent on which tissues the vessels supply (33, 52, 53) and the pregnant versus nonpregnant state of female animals (54). As such, it is imperative that the involvement of NO in MNP-induced uterine endothelial dysfunction in pregnancy is further explored. Elucidating how MNPs influence maternal blood flow to the fetoplacental compartment will inform expectations for the developmental consequences of this emerging contaminant. We hypothesize that MNP inhalation will negatively impact NO signaling in the uterine vasculature. Therefore, we aimed to examine the potential mechanisms of impaired NO signaling in uterine endothelial dysfunction after repeated MNP inhalation throughout pregnancy.

## MATERIALS AND METHODS

### Animals

Time-pregnant rats were ordered from Charles River Laboratories (Kingston, NY, USA) and housed in a Rutgers University AAALAC accredited vivarium. Normal chow and water were available to the rats *ad libitum*. Rats were given a minimum of 24 hours to acclimate prior to handling. Animals were randomly divided into control and polyamide MNP-exposed groups. All experiments were conducted with Rutgers University IACUC approval.

### Inhalation Exposure

Polyamide-12 MNPs (Orgasol® 2001 UD NAT 2, Arkema, King of Prussia, PA) were aerosolized via acoustic generator to reach a target concentration of 10.15 ± 1.36 mg/m^3^ within a whole-body exposure chamber as previously described (19, 55). Rats were exposed to MNPs via whole-body inhalation for 4h/day 5d/week from gestational day 5 to gestational day 19. Sampling was done in real time to characterize the size distribution of the aerosol using a Scanning Mobility Particle Sizer (SMPS, TSI Model 3080, Shoreview, MN) and High resolution electrical low-pressure impactor (HR-ELPI+; Dekati, Kangasala, Finland). Representative size characterization data can be found in Figure S1.

### Wire Myography

Vessel reactivity of the uterine artery, the major conduit artery of the uterus, after polyamide-12 exposure was assessed via wire myography (DMT, Ann Arbor, MI) as previously described (19, 20, 56). In brief, the uterine artery was excised and placed in cold Wire Myography Physiological Salt Solution (WM-PSS: 130 mM NaCl, 4.7 mM KCl, 1.18 mM KH_2_PO_4_, 1.17 mM MgSO4 7H_2_O, 14.9 mM NaHCO_3_, 5.5 mM glucose, 0.03 mM EGTA, and 1.6 mM CaCl_2_.Segments of the vessel (2 mm in length) were cut, mounted on two wires within 1 h of tissue harvest, and used for experimentation. Endothelium-dependent relaxation, endothelium-independent relaxation, and smooth muscle contraction were separately evaluated via cumulative additions of 60 µL of Methacholine chloride (MCh, MP Biomedicals, CAT 190231), Sodium Nitroprusside Dihydrate (SNP, Sigma-Aldrich, CAT 567538), and Phenylephrine (PE, Thermofisher Scientific, CAT 207240100), respectively for a concentration-response curve spanning from 10^− 9^ to 10^− 4^ M for each pharmacological application. Responses to chemical agents were randomized for each vessel segment. Uterine artery wire myography data are presented as a percentage of maximum tension after incubation with High Potassium Physiological Salt Solution (KPSS: 74.7 mM NaCl, 60 mM KCl, 1.18 mM KH_2_PO_4_, 1.17 mM MgSO_4_ 7H_2_O, 1.6 mM CaCl_2_, 14.9 mM NaHCO_3_, 0.03 mM EDTA, and 5.5 mM glucose) for 5 minutes, which reflects the maximum tension placed on the wire in the myograph chamber by the vessel. Tension on the wire at baseline and with the addition of the pharmacological agents was recorded and the percentage of maximum tension generation was calculated using the formula below:

Percentage maximum tension (%) = [(T_AGONIST_ −T_BASELINE_)/ T_MAX_] ×100, where T_AGONIST_ is the steady-state tension (mN) achieved after each chemical bolus, T_BASELINE_ is the control tension (mN) measured immediately prior to the dose-response experiment, and T_MAX_ is the maximal tension (mN) recorded after the addition of KPSS.

### Pressure Myography

Vessel reactivity of the radial artery, the resistance artery responsible for the perfusion of the placenta and a representation of the uterine microciruclation, after polyamide-12 exposure was assessed via pressure myography as previously described (19, 34). The uterine horn was placed in a bath of 4° C Microvessel Physiological Salt Solution (MV-PSS: 119 mM NaCl, 4.7 mM KCl, 1.17 mM MgSO_4_ 7H_2_O, 1.6 mM CaCl_2_ 2 H_2_O, 1.18 mM NaH_2_PO_4_, 24 mM NaHCO_3_, 5.5 mM glucose, and 0.03 mM EGTA) and a single radial artery was excised, mounted, and pressurized to 60 mmHg in a bath of MV-PSS before experimentation and within 1 h of tissue harvest. The equilibrated artery was incubated with various pharmacological agents (MCh, SNP, and PE) to measure concentration-dependent (10^− 9^ to 10^− 4^ M) physiological responses.

Pressure myography in radial artery segments was used to assess how modulators of eNOS function influence MCh-induced endothelium-dependent dilation in vessel segments from MNP-exposed dams. Responses of each radial artery segment to MCh (10^− 9^ to 10^− 4^ M) were recorded. Then, each vessel was incubated for 15 minutes with a randomly selected modulator of eNOS function. NG-Nitro-L-Arginine Methyl Ester, Hydrochloride (L-NAME, 100 µM, Sigma-Aldrich, CAT 483125), a nitric oxide synthase inhibitor, was used to determine the contribution of nitric oxide synthases to MCh-induced vasodilation. The modulator L-Arginine (100 µM, Sigma-Aldrich, CAT A5006), an eNOS substrate was used to identify potential substrate deficiency between groups. The modulator, tetrahydrobiopterin (BH_4_, 100 µM, Thermofisher Scientific, CAT 328871000), an eNOS cofactor, was used to identify cofactor deficiency. Dithiothreitol (DTT, 300 nM, Thermofisher Scientific, CAT R0861), a reducing agent, was used to reverse oxidative modifications to eNOS that modulate function. After incubation with the selected modulator, the concentration-dependent response of each radial artery segment to MCh (10^− 9^ to 10^− 4^ M) was measured.

Pressure myography data are presented as a percentage of maximum relaxation after incubation with Ca^2+^ Free Microvessel Physiological Salt Solution (Ca^2+^ Free PSS: 120.6 mM NaCl, 4.7 mM KCl, 1.17 mM MgSO_4_ 7H_2_O, 1.18 mM NaH_2_PO_4_, 24 mM NaHCO_3_, 5.5 Mm glucose, and 0.03 mM EGTA) for 20 minutes, which reflects the maximum dilation of the vessel measured by the caliper when maximum dilation was elicited by Ca^2+^ Free PSS. Pressure myography data is presented as a percentage of maximum dilation which was quantified using the formula below:

Percentage maximum dilation (%) = [(D_POST_−D_BASELINE_)/(D_MAX_−D_BASELINE_)] × 100, where D_POST_ is the steady-state diameter (µm) achieved after each chemical bolus, D_BASELINE_ is the control diameter (µm) measured immediately prior to the dose-response experiment, and D_MAX_ is the maximal diameter (µm) recorded after the addition of Ca^2+^ Free PSS.

### Oxymyoglobin oxidation

Oxymyoglobin (OxyMb) oxidation by whole uterine vasculature was measured to determine MCh-stimulated tissue production of NO-related species. A 200 nM solution of native equine heart myoglobin (Sigma-Aldrich, CAT M1882) was produced and an absorbance spectrum between 476 and 700nm was generated (Figure S2A). The metmyoglobin (MetMb) was reduced with sodium dithionite to generate deoxymyoglobin (DeoxyMb), inducing a spectral change. After centrifugal filtration for 20 minutes at 4,000xg, the myoglobin was resuspended in PSS to generate 0.2mM oxymyoglobin (OxyMb). A spectrum was generated for the OxyMb (Figure S2B) and an isosbestic point for Native Mb and OxyMb was identified at 594nm (Figure S2C). OxyMb was warmed to 37° C for 45 minutes and spectra were generated for the warmed OxyMb. Next, fresh pooled uterine vessels were dried of excess fluid, added to the warmed OxyMb solution, and allowed 5 minutes to equilibrate to 37° C after which a spectrum was generated. MCh was added to the solution for a final concentration of 10^-4^ M to elicit NO production. Spectra were generated at 3 and 15 minutes after the addition of MCh. The tissue was then rinsed of Mb and snap frozen on dry ice.

Estimated production of NO-related species was based on the difference spectra of native Mb and OxyMb at absorbance values at wavelengths of 580nm (point with largest absolute value) and 594nm (isosbestic point). This difference spectrum represents the maximum possible oxidation of OxyMb (Figure S2D). Experimental oxidation of OxyMb was determined using a difference spectrum of experimental Mb (at 3 or 15 minutes) and native Mb. Absorbance values at 580nm and 594nm represent the experimental oxidation of OxyMb. OxyMb oxidation (nmoles) was calculated as a proportion of the maximum possible oxidation using the formula below and normalized to tissue weight.

nmoles oxidized OxyMb= [(Experimental Difference Absorbance_594_−Experimental Difference Absorbance_580_)/(Maximum Oxidation Difference Absorbance_594_− Maximum Oxidation Difference Absorbance_580_)] × 200nM

### Whole Uterine Vasculature Protein Isolation

Samples were crushed using the cell crusher homogenization tool at frozen temperatures and added to RIPA buffer with protease and phosphatase inhibitors (Thermofisher Scientific CAT 89900 and 78442). After a 30-minute digestion at 4° C, the mixture was then centrifuged at 14,000 RPM at 4° C for 25 minutes. The supernatant was then isolated and centrifuged at 14,000 RPM at 4° C for 15 minutes. The supernatant was removed and assessed for protein concentration using the Pierce™ BCA Protein Assay Kit (Thermofisher Scientific CAT 23227).

### Western Blot

Isolated protein (30 µg) was run on a 4-12% bis tris gel (Thermofisher Scientific CAT WG1402BOX). Proteins were transferred onto a PVDF membrane using the Biorad Trans-Blot Turbo system (CAT 1704150). Membranes were blocked with 5% skim milk in TBST and probed with primary antibodies against eNOS, phospho-eNOS Ser^1176^ (Ser1177 in humans), arginase-1, GTPCH, DHFR, 3-nitrotyrosine residues, Thioredoxin, GCLC, and β-actin. Membranes were incubated with anti-rabbit or anti-mouse HRP-conjugated secondary antibodies and developed using SuperSignal West Dura substrate (Thermofisher Scientific CAT 34075). Band intensity was quantified using Image J. Protein expression was normalized to β-actin. Data are presented with respect to fold change of protein expression relative to control samples. Antibody sources and concentrations can be found in Table S1.

### Statistics

Wire myography and pressure myography data were analyzed by nonlinear regression analysis. Maternal/litter characteristics and Western blot data were analyzed by a two-tailed t-test assuming equal variance between groups or Welch’s t-test when variance was significantly different between groups. Data are expressed as mean ± standard error with the dam/litter as the sample size unit. Statistical analyses were completed with GraphPad Prism 9.0 (San Diego, CA).

## RESULTS

### Maternal and Litter Characteristics

MNP inhalation during pregnancy led to a modest reduction in maternal weight, although, this was not statistically significant (Table 1, p = 0.09). Maternal mean arterial pressure was unchanged after exposure (Table 1). Fetal weight, however, was significantly reduced and placental weight was significantly increased in exposed litters. Placental efficiency, grams of pup weight produced per gram of placental tissue, a measure of placental function, was diminished in MNP-exposed litters (Table 1). Together, these data show that the developing offspring are susceptible to the effects of maternal MNP inhalation.

**Table 1.** Maternal and litter characteristics of control and polyamide-exposed Sprague Dawley rats.

| Measurement | Control | Polyamide |
| --- | --- | --- |
| Maternal Weight (g) | 312 ± 3 (n=57) | 301 ± 6 (n=32) |
| Maternal Mean Arterial Pressure (mmHg) | 73.8 ± 1.4 (n=30) | 72.6 ± 2.0 (n=22) |
| Fetal Weight (g) | 4.41 ± 0.03 (n=52) | 4.23 ± 0.08 (n=31)* |
| Placental Weight (g) | 0.49 ± 0.01 (n=52) | 0.52 ± 0.01 (n=30)* |
| Placental Efficiency (fetal weight/placental weight) | 9.03 ± 0.13 (n=52) | 8.19 ± 0.21 (n=31)* |
| Resorptions | 0.42 ± 0.08 (n=56) | 0.50 ± 0.13 (n=32) |
| Litter Size | 9.62 ± 0.32 (n=50) | 9.41 ± 0.41 (n=32) |

Values are presented as mean ± SEM with the dam/litter as the statistical unit of comparison. Significant differences ( $p < 0.05$ ) from the control group are denoted by \*.

### Uterine Artery Vascular Responsivity

Responsivity to stimuli in the uterine artery, the major conduit artery of the uterus, was assessed *ex* vivo via wire myography in control and MNP-exposed dams (Figure 1). No significant differences in reactivity were noted in endothelium-dependent relaxation (Figure 1A), endothelium-independent relaxation (Figure 1B), and smooth muscle contraction (Figure 1C). These results demonstrate that MNP inhalation throughout pregnancy does not impair uterine artery vascular reactivity.

**Figure 1.**
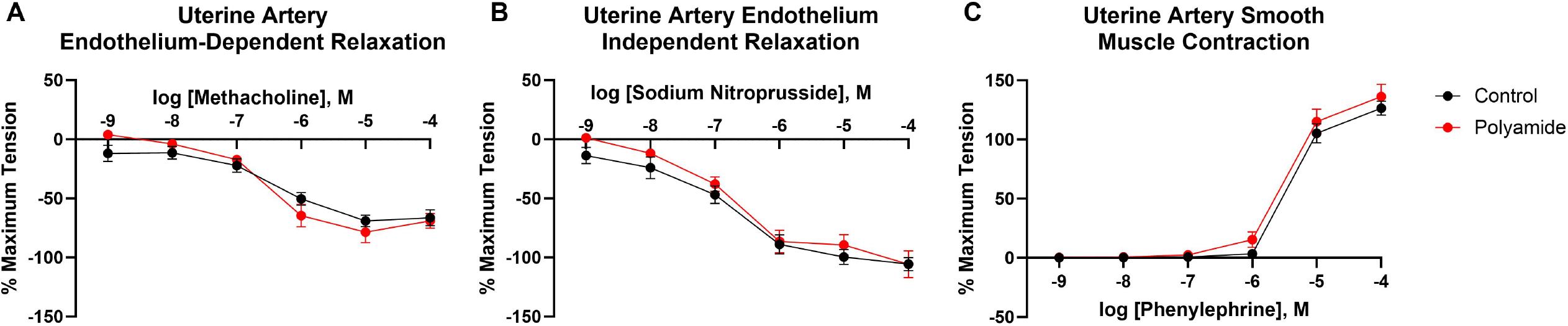
Vascular reactivity of the uterine artery was assessed via wire myography. Concentration-response curves were generated after treatment of uterine artery segments with increasing concentrations of the endothelial-dependent vasodilator methacholine (A), the endothelial-independent vasodilator sodium nitroprusside (B), and the vasoconstrictor phenylephrine (C). Significance was assessed by comparing overall reactivity via a four-parameter nonlinear regression analysis. Data are presented as mean ± SEM, n =7-10.

### Radial Artery Vascular Responsivity

*Ex vivo* function of radial arteries, the major resistance vessels of the uterus that regulate placental perfusion, were assessed via pressure myography (Figure 2). No anatomical changes in radial arteries were observed between groups (Table S2). MNP exposure significantly blunted endothelium-dependent dilation of the radial artery (Figure 2A) while endothelium-independent dilation was unchanged (Figure 2B). Smooth muscle constriction in response to PE was not different between groups (Figure 3C). L-NAME inhibition of nitric oxide synthases abolished endothelium-dependent dilation in control and exposed radial arteries, demonstrating the reliance of both groups on nitric oxide synthase signaling (Figure 2D). Overall, MNP exposure induced endothelial dysfunction and had no effect on vascular smooth muscle activity within the radial artery of the uterus.

**Figure 2.**
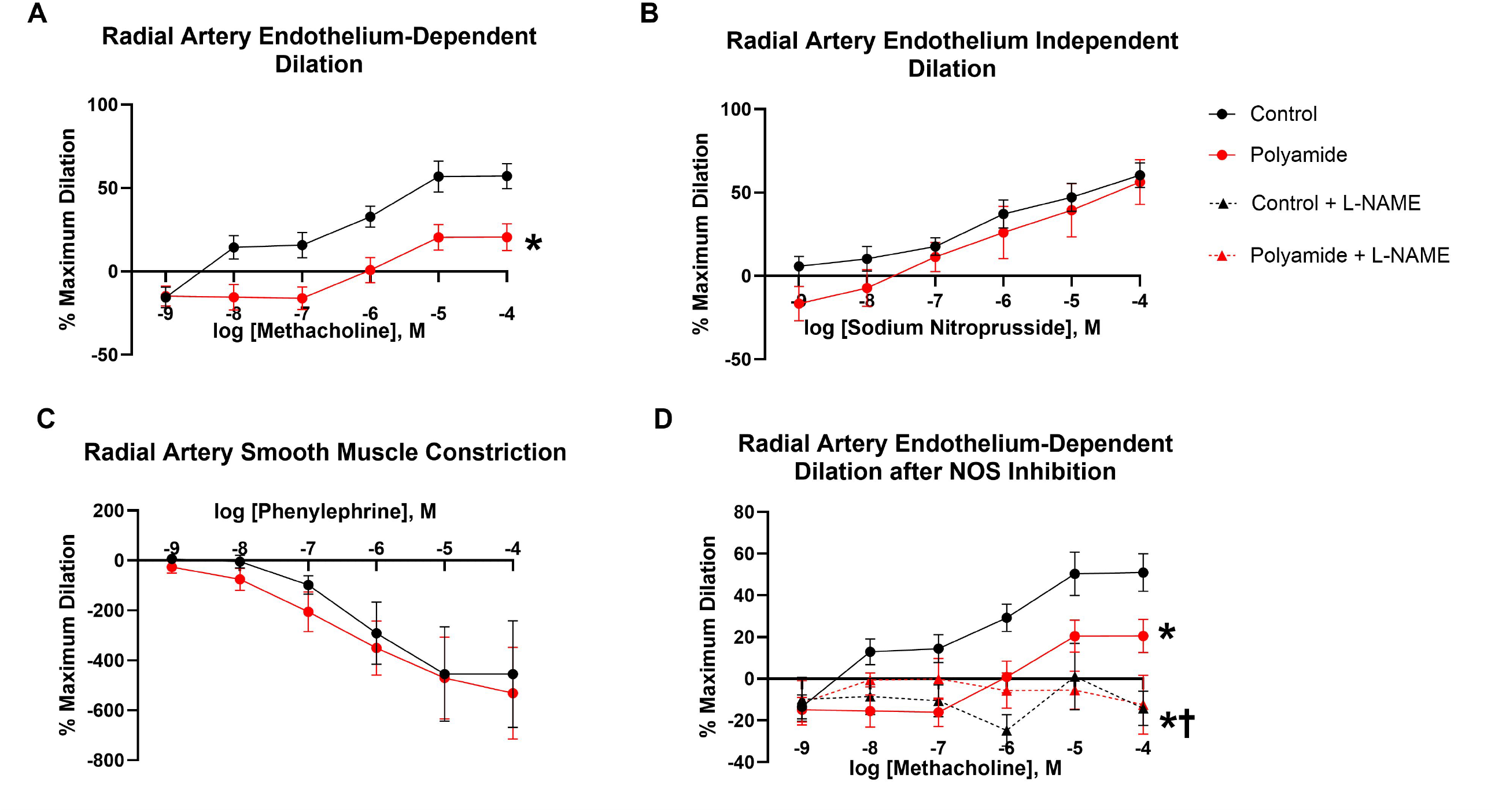
Vascular reactivity of the radial artery was assessed via pressure myography. Concentration-response curves were generated after treatment of radial artery segments with increasing concentrations of the endothelial-dependent vasodilator methacholine (A), the endothelial-independent vasodilator sodium nitroprusside (B), and the vasoconstrictor phenylephrine (C). Significance was assessed by comparing overall reactivity via a four-parameter nonlinear regression analysis. Data are presented as mean ± SEM, n =8-10. Significant differences (p < 0.05) from the control group are denoted by * and significant differences from the polyamide-exposed group are denoted by †.

**Figure 3.**
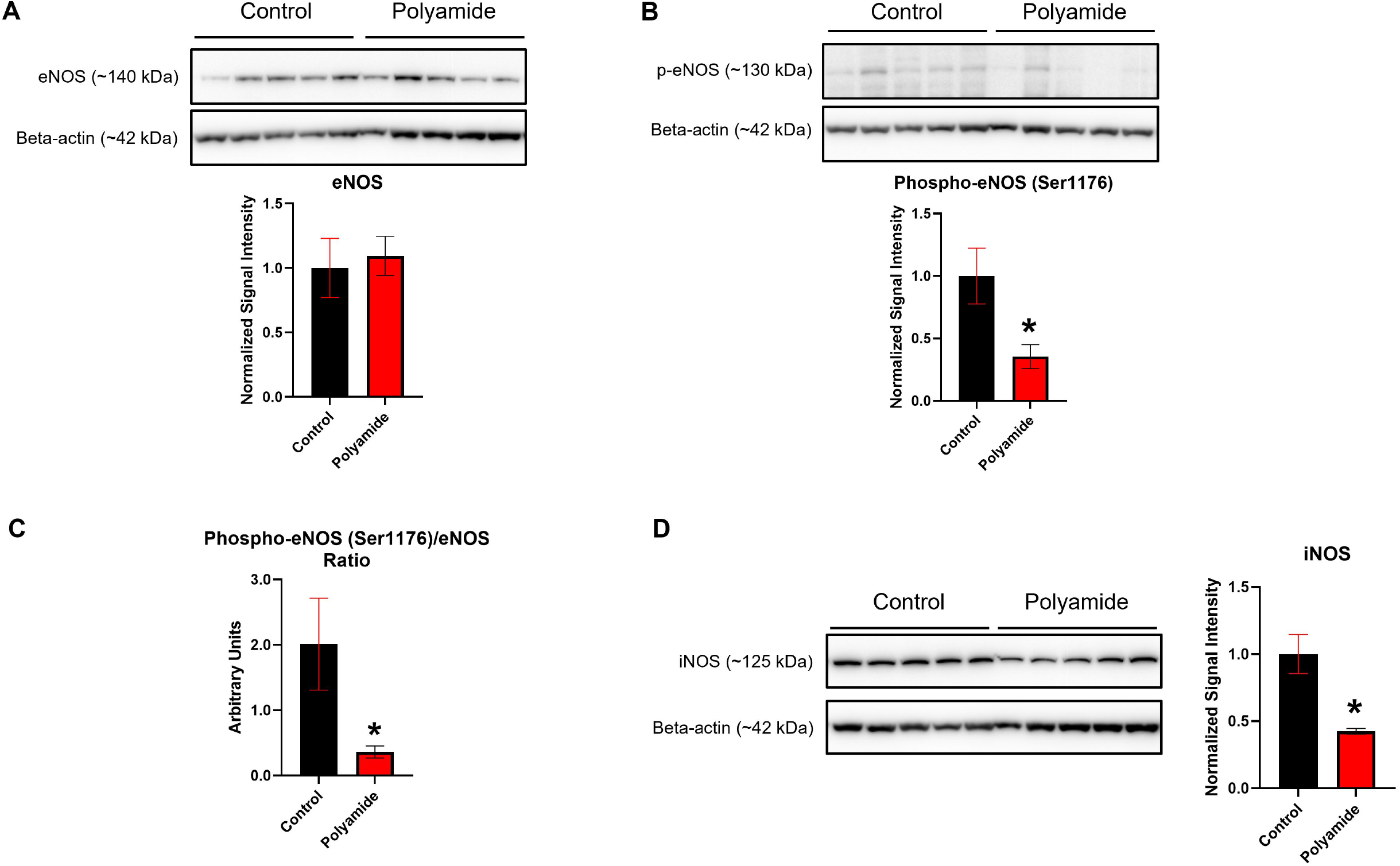
Western Blot of uterine vasculature from control and MNP-exposed dams was carried out to determine protein expression of eNOS (A), phospho-eNOS (Ser1176) (B), the ratio of phospho-eNOS to eNOS (C), and iNOS (D). Significance was assessed through two-tailed Student’s t-test. Data are presented as mean ± SEM, n =9. Significant differences (p < 0.05) between the control and polyamide-exposed groups are denoted by *.

### Nitric Oxide Synthase Expression in Uterine Vasculature

Protein expression of eNOS and inducible nitric oxide synthase (iNOS) were measured in the uterine vascular bed of control and MNP-exposed dams (Figure 3). There was no significant difference in eNOS expression between groups (Figure 3A). However, relative levels of phospho-eNOS (Ser1176), a post-translational modification of eNOS that facilitates electron flow through eNOS, was significantly decreased in uterine vascular beds of exposed dams (Figure 3B). The ratio of phospho-eNOS to eNOS was significantly lower in exposed dams which suggests that MNP exposure reduced the amount of activated eNOS in the uterine vascular bed (Figure 3C). Inducible NOS (iNOS) protein expression was assessed as this NOS may contribute to NOS activity. Interestingly, iNOS expression was significantly lower in uterine vasculature of MNP-exposed dams (Figure 4D). Taken together, these data suggest that eNOS activation is decreased and iNOS is not induced by exposure.

**Figure 4.**
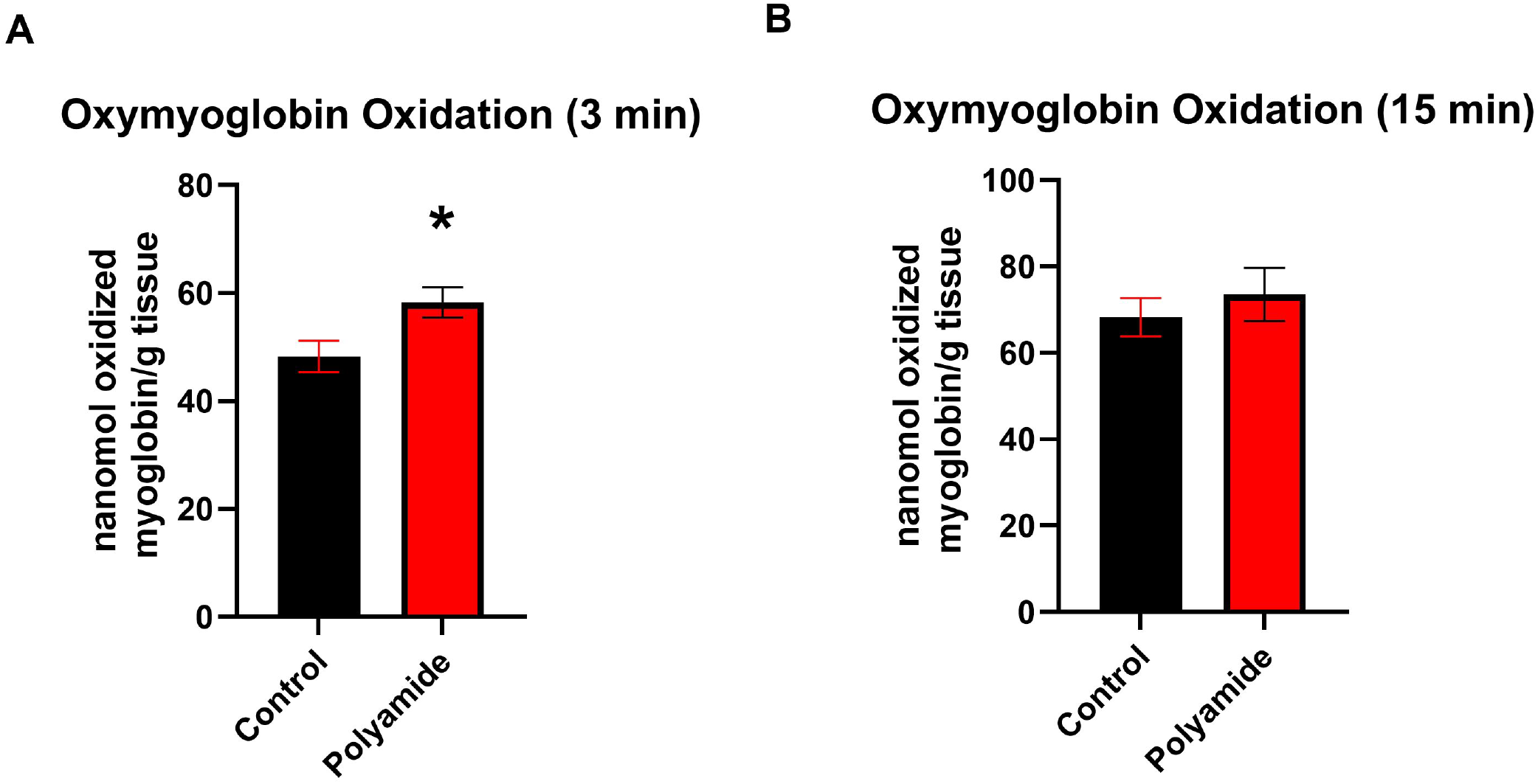
Myoglobin oxidation by live uterine vasculature was measured as a surrogate for NO-related species production. Oxymyoglobin oxidation was measured at 3 minutes (A) and 15 minutes (B) after stimulation of vascular tissue with methacholine. Significance was assessed through two-tailed Student’s T-Test. Data are presented as mean ± SEM, n =9. Significant differences (p < 0.05) between the control and polyamide-exposed groups are denoted by *.

### OxyMb Oxidation by Uterine Vascular bed

OxyMb oxidation was determined as a representative measure of NO derived from the uterine vascular beds of control dams and dams exposed to MNP via inhalation (Figure 4). Initially, the uterine vasculature from exposed dams oxidized more moles of OxyMb per mg tissue than controls after stimulation with 10^-4^ MCh (3 minutes) (Figure 4A). However, by 15 minutes, oxidation of OxyMb was not significantly different (Figure 4B). These data suggest that the vasculature of polyamide-exposed dams experiences a short-term burst of production of NO after stimulation by MCh.

### L-Arginine intervention of MNP-induced endothelial dysfunction

The role of L-Arginine, the enzymatic substrate of eNOS, in MNP-induced endothelial dysfunction was investigated (Figure 5). Endothelium-dependent dilation of radial artery segments from MNP-exposed dams was not improved after *ex vivo* L-Arginine supplementation (Figure 5A). Relative expression of arginase 1, which catalyzes the conversion of L-Arginine to ornithine, was not different between groups (Figure 5B). These data demonstrate that there is no L-Arginine deficiency in MNP-exposed uterine vasculature.

**Figure 5.**
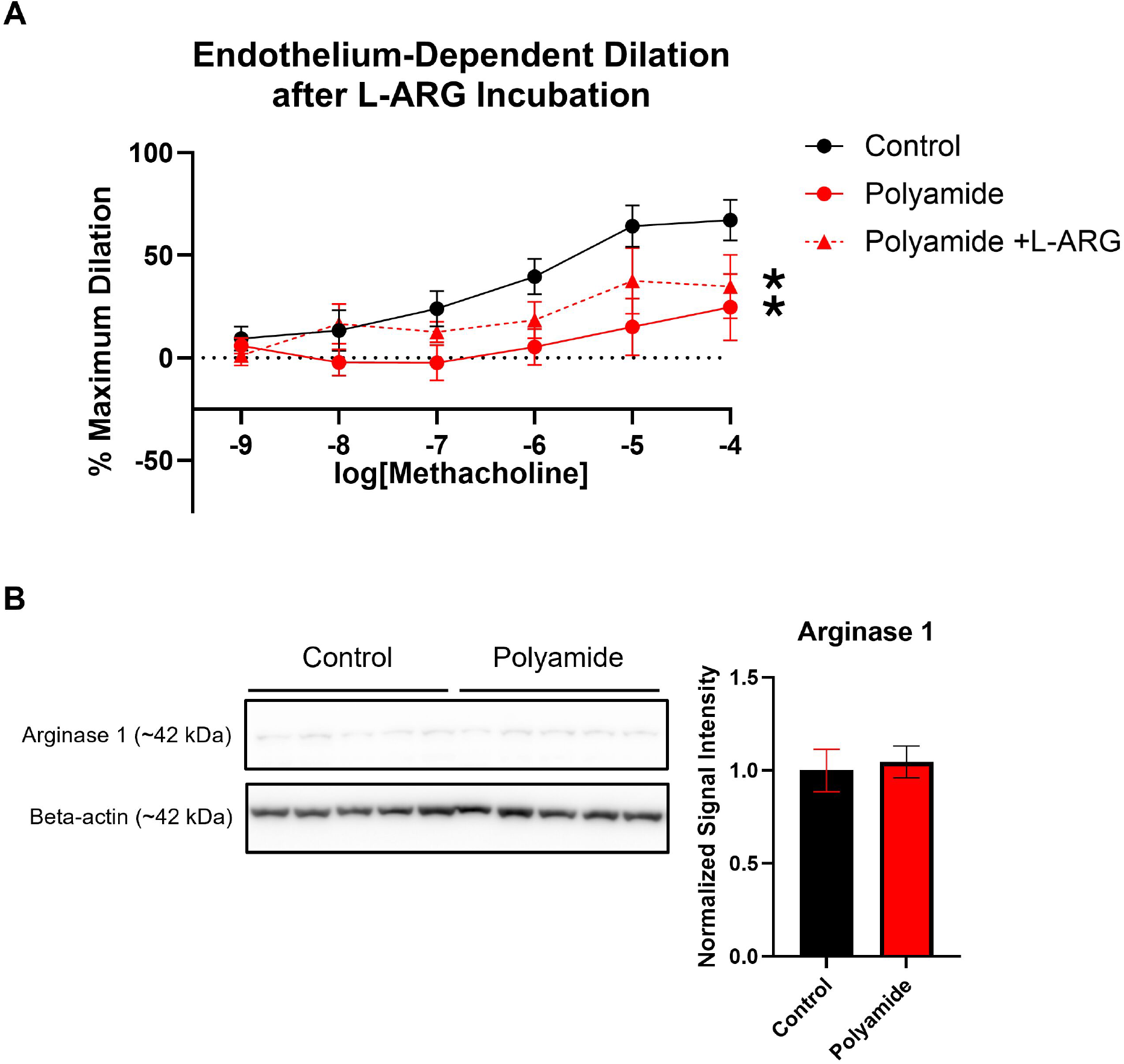
The effect of L-arginine supplementation on radial endothelium-dependent vasodilation and arginase protein expression in whole uterine vasculature was assessed. After incubation of radial artery segments with L-arginine, concentration-response curves were generated via treatment of segments with increasing concentrations of the endothelial-dependent vasodilator methacholine (A). Relative protein expression of arginase 1 in uterine vasculature was determined by Western Blot (B). Significance was assessed by comparing overall reactivity via a four-parameter nonlinear regression or two-tailed Student’s t-test analysis for vascular reactivity and Western Blot, respectively. Data are presented as mean ± SEM, n =8-9. Significant differences (p < 0.05) from the control group is denoted by *.

### BH_4_ rescue of endothelial dysfunction

The influence of BH_4_, a cofactor of eNOS was investigated. BH_4_ supplementation significantly improved endothelial-dependent dilation in MNP-exposed radial artery segments, although dilation was not restored to control levels (Figure 6A). The expression of GTP Cyclohydrolase (GTPCH), the rate-limiting enzyme in the de novo production of BH_4_ (57), was significantly increased in MNP-exposed dams (Figure 6B). Dihydrofolate Reductase (DHFR), an enzyme responsible for (dihydrobiopterin) BH_2_ reduction to BH_4_ (57), revealed an increase in expression after MNP exposure (p = 0.09), although this was not statistically significant (Figure 6C). These data suggest that eNOS within the uterine vasculature of MNP-exposed dams may be uncoupled due to a BH_4_ deficiency despite increased cofactor production.

**Figure 6.**
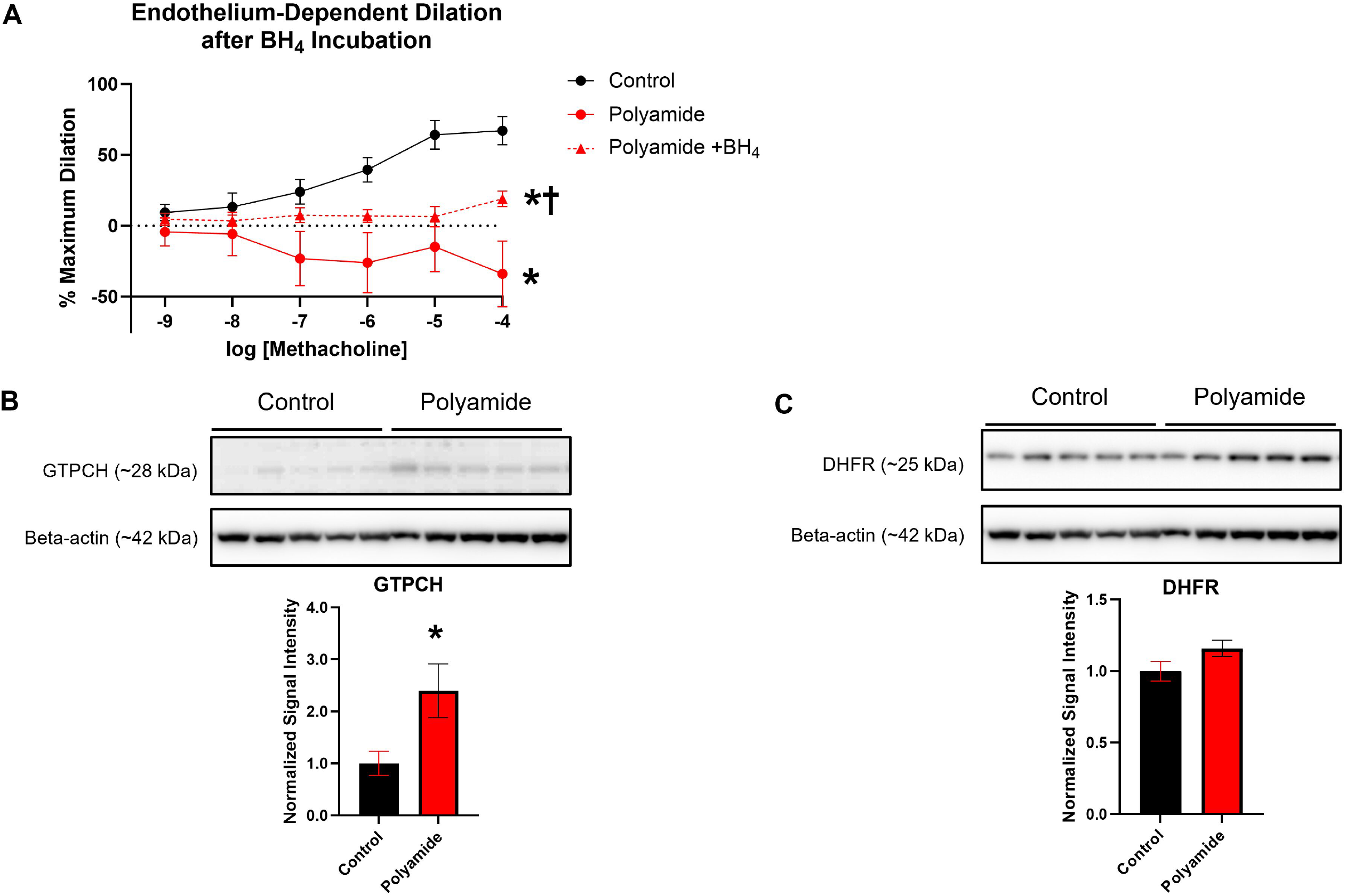
The effect of BH_4_ supplementation on radial endothelium-dependent vasodilation and GTPCH and DHFR protein expression in whole uterine vasculature was assessed. After incubation of radial artery segments with BH_4_, concentration-response curves were generated via treatment of segments with increasing concentrations of the endothelial-dependent vasodilator methacholine (A). Relative protein expression of GTPCH (B) and DHFR (C) in uterine vasculature was determined by Western Blot. Significance was assessed by comparing overall reactivity via a four-parameter nonlinear regression or two-tailed Student’s t-test analysis for vascular reactivity and Western Blot, respectively. Data are presented as mean ± SEM, n =8-9. Significant differences (p < 0.05) from the control group are denoted by *. Significant differences (p < 0.05) from the polyamide-exposed group are denoted by †.

### Contribution of oxidative/nitrosative stress to endothelial dysfunction

The role of oxidative/nitrosative stress on MNP-induced endothelial dysfunction in radial arteries was examined (Figure 7). Incubation of radial arteries from MNP-exposed dams with the reducing agent DTT significantly improved endothelium-dependent dilation, although vasodilation function was not restored to control levels (Figure 7A). Nitration of tyrosine residues, a product of peroxynitrite formation, was increased in the uterine vasculature after MNP exposure during pregnancy (Figure 7B). Relative levels of thioredoxin, an antioxidant enzyme that reverses oxidative modifications to eNOS, were significantly reduced after MNP exposure (Figure 7C). In the uterine vascular bed of exposed dams, glutamate cysteine ligase catalytic subunit (GCLC), the rate-limiting enzyme in glutathione production (58), was comparable to controls (Figure 7D). Together, these data suggest MNP inhalation promotes an oxidative environment in the uterine vasculature, disrupting electron flow through eNOS, preventing NO production, and ultimately impairing endothelial function in the radial artery.

**Figure 7.**
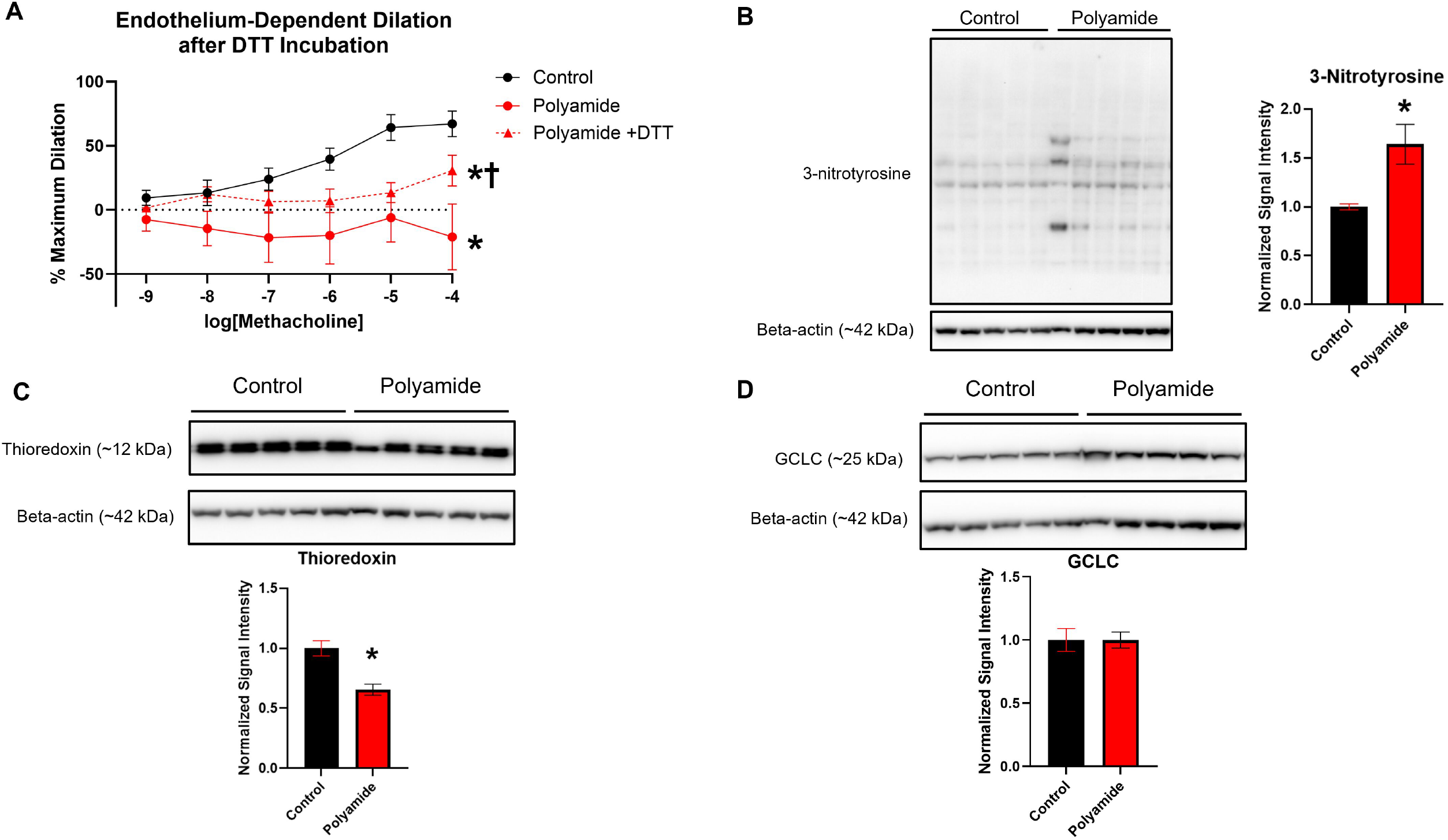
The effect of DTT on radial endothelium-dependent vasodilation function and 3-nitrotyrosine, thioredoxin, and GCLC protein expression in whole uterine vasculature was assessed. After incubation of radial artery segments with DTT, concentration-response curves were generated via treatment of segments with increasing concentrations of the endothelial-dependent vasodilator methacholine (A). Relative protein expression of 3-nitrotyrosine residues (B) thioredoxin (C) and GCLC (D) in uterine vasculature was determined by Western Blot. Significance was assessed by comparing overall reactivity via a four-parameter nonlinear regression or two-tailed Student’s t-test analysis for vascular reactivity and Western Blot, respectively. Data are presented as mean ± SEM, n =8-9. Significant differences (p < 0.05) from the control group are denoted by *. Significant differences (p < 0.05) from the polyamide-exposed group are denoted by †.

## DISCUSSION

### Major findings and implications

Herein, we found that repeated MNP inhalation exposure during pregnancy induced endothelial dysfunction in the uterine microcirculation but not conduit vessels. This disparity in vascular dysfunction between conduit vessels and resistance vessels has been observed previously after a pulmonary MNP exposure in late gestation (20). Abrogated endothelial function reduces vasodilation of the uterine microcirculation and limits the ability of the microcirculation to meet the metabolic demand of the fetoplacental unit. Therefore, MNP-induced uterine endothelial dysfunction may account for the fetal growth restriction we observed in our study (Table 1). Moreover, limited perfusion of the fetoplacental unit generates a hostile intrauterine environment and adversely impacts offspring cardiometabolic health (59-61). Through nitric oxide synthase inhibition, we demonstrated NO signaling is crucial for endothelial-dependent dilation in the radial artery of control and MNP-exposed dams which is consistent with other studies of radial artery signaling (62). We noted that MNP-exposed uterine vascular tissue led to significantly heightened OxyMb oxidation and, therefore, more production of NO than control tissue. Considering the decreased eNOS activation and iNOS expression, it is likely that the observed increase in OxyMb oxidation is due to elevated NO-related species that do not elicit dilation in the uterine vasculature such as peroxynitrite, the reaction product of superoxide and NO, or nitrites (63-65). This is also consistent with the identified reduction in nitric oxide bioavailability and increase of other NO-related species like peroxynitrite or nitrites after nano-TiO_2_ and ultrafine PM exposure (48, 66, 67). Taken together, these data support that exposure impaired NO signaling, leading to endothelial dysfunction, blunted vasodilation, and reduced resources for fetal growth. Critically this may inhibit metabolic perfusion matching in the placenta which could seriously limit oxygen-dependent metabolism.

### Uncoupling and deactivation of eNOS as mechanisms for uterine MNP-induced microvascular vascular toxicity

The physiological utility of eNOS is reliant on electron flow from the reductase to the oxygenase domain within the homodimer. To produce NO from L-arginine and molecular oxygen, electrons must travel from NADPH through FAD, FMN, BH_4_, and ultimately to L-arginine. When this electron transfer is disrupted, eNOS cannot direct the electrons to L-arginine and instead produces superoxide molecules. In this state, the transfer of electrons from NADPH is no longer coupled to NO production; therefore, eNOS is described as “uncoupled” as observed in diabetes, hypercholesterolemia, and hypertension (68). Notably, BH_4_ deficiency is one of the most frequently identified causes of eNOS uncoupling in human disease and experimental disease models (69). In addition to coupling, another determinant of eNOS function are post-translational modifications, several of which can either enhance or dampen NO production (70).

Activating modifications, including phosphorylation at serine 1176 (1177 for humans) on eNOS, facilitate faster electron transfer in the reductase domain and are thought to activate eNOS, increasing NO bioavailability (71).

In our study, the prevalence of the activating post-translational modification, Ser^1176^, on eNOS was decreased by over 50% in exposed uterine vasculature. Therefore, it can be deduced that MNP exposure decreases the activation of eNOS in the uterine vasculature. PM_2.5_ has similarly decreased eNOS activation in mice (72). Improvement of radial endothelium-dependent dilation in MNP-exposed tissue was achieved with the addition of BH_4_, demonstrating a deficiency of this cofactor which is consistent with other models of PM exposure (50, 51, 73). Interestingly, elevated GTPCH1 expression in the uterine vasculature of exposed dams suggests that BH_4_ production is increased. This contrasts with studies showing that cigarette smoke exposure increases GTPCH1 protein expression (73) or that PM_2.5_ has no apparent effect on GTPCH1 expression (74). The evident deficiency and increased BH_4_ production in our model suggest that an external factor depletes BH_4_ in the tissue, resulting in eNOS uncoupling. In the MNP-exposed group, improvement of radial artery function by the reducing agent DTT, decreased thioredoxin levels, and elevated 3-nitrotyrosine residues in tissue demonstrate an oxidative or nitrosative environment. Other particle exposures similarly produce oxidative and nitrosative species in vascular tissue (74). This environment introduces deactivating post-translational modifications that promote eNOS uncoupling (75). In turn, eNOS produces superoxide ions instead of NO, further exacerbating the nitrosative and oxidative environment, depleting BH_4_, and ultimately promoting more eNOS dysfunction (51, 76). Therefore, our data suggests repeated MNP inhalation uncouples eNOS, which is consistent with vascular studies of pulmonary particle exposure (47-51).

### Study limitations

While this study is the first to clarify the role of NO signaling in MNP-induced endothelial dysfunction, some aspects of our study design limit the potential extrapolation of our results. The investigations in this study were carried out in pregnant dams. Because successful pregnancy requires hormonal changes and physiological adaptations, including an increase in systemic estradiol, which supports vasodilation, different outcomes may occur in nonpregnant female or male rats (54, 77). The uterine vasculature was the only vascular bed assessed in our study and has distinct responses to stimuli when compared to vessels perfusing other organs (78). Given that particle exposure can affect each vascular bed differently (52), outcomes in the coronary, mesentery, or skeletal vasculature may differ from our findings in the uterine vasculature. As stated previously, multiple signaling pathways are enhanced during pregnancy to promote uterine vasodilation, including reliance on estrogen signaling (79-81). Previous work in our laboratory identified impaired uterine endothelial-dependent dilation and a significant reduction in circulating 17β-estradiol in nonpregnant rats after a single inhalation exposure to polyamide-12 aerosols during estrus (19). Of the vascular interventions presented in the study, neither BH_4_ nor DTT fully restored function to control levels, supporting the involvement of other compensatory mechanisms of endothelial dysfunction. While NO signaling contributes to more than half of uterine vasodilation (40), prostacyclin production and intracellular calcium signaling may be impaired by MNP exposure or activated as a compensatory mechanism. The role of endogenous eNOS uncouplers, including asymmetric dimethylarginine, was not assessed in this study and may also contribute to the observed eNOS dysfunction.

The polyamide-12 polymer used in this study has unique physicochemical properties due to its chemical structure and spheroidal shape. Plastics have the ability to adsorb chemicals to their surface and release impregnated plasticizing chemicals or oxidative molecules into the local area. We have identified the translocation of plastic particles from the maternal lungs to and through the placenta, depositing within the fetal and remaining in the F1 generation tissues (55, 82). These findings indicate the plausibility of direct particle interactions which may dictate endothelial cell health and function. These unique properties influence interactions with tissue and overall toxicity. Therefore, other plastic polymers, including polystyrene, polyvinyl chloride, or polyethylene may generate different vascular toxicities. Likewise, the MNPs used in our investigation were considered naïve/pristine particles that did not undergo weathering processes such as UV oxidation, mechanical abrasion, or thermal degradation.

### Future mechanistic studies and considerations

In this study, no tested intervention completely restored endothelial-dependent dilation to levels seen in control radial arteries. Therefore, the observed endothelial dysfunction may involve other mechanisms that influence vasodilation. For example, calcium ion mobilization in the endothelial cell after MCh stimulation may be impaired, leading to decreased eNOS activation. The degree to which oxidative and nitrosative stress in tissue specifically affects eNOS function is unknown. Future studies identifying whether eNOS was selectively nitrated and if eNOS was nitrated at specific activating or deactivating phosphorylation sites would elucidate how MNPs generate uncoupling of eNOS. Furthermore, other enzymes, such as cyclooxygenases, may have been activated in the MNP-exposed vasculature to compensate for eNOS uncoupling. Subsequent studies should identify secondary mechanisms that may correct for loss of eNOS function.

### Conclusions

The reduction of uterine vascular endothelial function by MNPs suggests that MNP inhalation poses serious repercussions for fetal health. As plastic waste continues to enter our environment at an exponential pace, it should be expected that health complications will follow suit. As such, continued research on MNP toxicity should aim to identify mechanistic targets as this will aid in the development of early interventions for pregnant women. This study underscores eNOS deactivation, BH_4_ depletion, and nitrosative and oxidative stress as important factors in MNP-induced endothelial dysfunction. Therefore, dietary changes or supplementation to promote antioxidant capabilities and BH_4_ production may partially abrogate the vascular toxicity of MNPs.

## DATA AVAILABILITY

Source data for this study are not publicly available due to privacy restrictions. The source data are available upon request by contacting the corresponding author.

## SUPPLEMENTAL MATERIAL

Supplemental Figure S1 https://doi.org/10.7910/DVN/NA1QL1

Supplemental Figure S2 https://doi.org/10.7910/DVN/LCYJQK

Supplemental Table S1 https://doi.org/10.7910/DVN/CJQULX

Supplemental Table S2 https://doi.org/10.7910/DVN/8BAW0L

## ACKNOWLEDGMENTS

We would like to thank Tanisha Brunson-Malone for facilitating the whole-body aerosol exposure and exposure sampling.

## AUTHOR CONTRIBUTIONS

Chelsea M. Cary, Andrew J. Gow, and Phoebe A. Stapleton conceived and designed the research included in this study. Chelsea M. Cary and Taina L. Moore performed experiments presented in this study. Chelsea M. Cary, Taina L. Moore and Phoebe A. Stapleton analyzed the data utilized in this manuscript. Chelsea M. Cary, Taina L. Moore, Andrew J. Gow, and Phoebe A. Stapleton interpreted results of the present experiments. Chelsea M. Cary prepared the present figures. Chelsea M. Cary drafted the manuscript. Chelsea M. Cary, Taina L. Moore, Andrew J. Gow, and Phoebe A. Stapleton edited and revised the manuscript. Phoebe A. Stapleton approved final version of the manuscript.

## REFERENCES

1. Torres-Agullo A, Karanasiou A, Moreno T, and Lacorte S. Airborne microplastic particle concentrations and characterization in indoor urban microenvironments. Environmental Pollution 308: 119707, 2022.

2. Xie Y, Li Y, Feng Y, Cheng W, and Wang Y. Inhalable microplastics prevails in air: Exploring the size detection limit. Environment International 162: 107151, 2022.

3. Sharaf Din K, Khokhar MF, Butt SI, Qadir A, and Younas F. Exploration of microplastic concentration in indoor and outdoor air samples: Morphological, polymeric, and elemental analysis. Science of The Total Environment 908: 168398, 2024.

4. Soltani NS, Taylor MP, and Wilson SP. Quantification and exposure assessment of microplastics in Australian indoor house dust. Environmental Pollution 283: 117064, 2021.

5. Liu C, Li J, Zhang Y, Wang L, Deng J, Gao Y, Yu L, Zhang J, and Sun H. Widespread distribution of PET and PC microplastics in dust in urban China and their estimated human exposure. Environment International 128: 116–124, 2019.

6. Jenner LC, Rotchell JM, Bennett RT, Cowen M, Tentzeris V, and Sadofsky LR. Detection of microplastics in human lung tissue using μFTIR spectroscopy. Sci Total Environ 831: 154907, 2022.

7. Amato-Lourenço LF, Carvalho-Oliveira R, Júnior GR, Dos Santos Galvão L, Ando RA, and Mauad T. Presence of airborne microplastics in human lung tissue. J Hazard Mater 416: 126124, 2021.

8. O’Neill MS, Veves A, Zanobetti A, Sarnat JA, Gold DR, Economides PA, Horton ES, and Schwartz J. Diabetes enhances vulnerability to particulate air pollution-associated impairment in vascular reactivity and endothelial function. Circulation 111: 2913–2920, 2005.

9. Jiang Y, Zhu X, Shen Y, He Y, Fan H, Xu X, Zhou L, Zhu Y, Xue X, Zhang Q, Du X, Zhang L, Zhang Y, Liu C, Niu Y, Cai J, Kan H, and Chen R. Mechanistic insights into cardiovascular effects of ultrafine particle exposure: A longitudinal panel study. Environ Int 187: 108714, 2024.

10. Zhang H, Yang J, Zhang Y, Xiao K, Wang Y, Si J, Li Y, Sun L, Sun J, Yi M, Chu X, and Li J. Age and sex differences in the effects of short- and long-term exposure to air pollution on endothelial dysfunction. Environ Health 23: 63, 2024.

11. de Bont J, Jaganathan S, Dahlquist M, Persson Å, Stafoggia M, and Ljungman P. Ambient air pollution and cardiovascular diseases: An umbrella review of systematic reviews and meta-analyses. J Intern Med 291: 779–800, 2022.

12. Khosravipour M, Safari-Faramani R, Rajati F, and Omidi F. The long-term effect of exposure to respirable particulate matter on the incidence of myocardial infarction: a systematic review and meta-analysis study. Environ Sci Pollut Res Int 29: 42347–42371, 2022.

13. Rundell KW, Hoffman JR, Caviston R, Bulbulian R, and Hollenbach AM. Inhalation of ultrafine and fine particulate matter disrupts systemic vascular function. Inhal Toxicol 19: 133–140, 2007.

14. Cuevas AK, Liberda EN, Gillespie PA, Allina J, and Chen LC. Inhaled nickel nanoparticles alter vascular reactivity in C57BL/6 mice. Inhal Toxicol 22 Suppl 2: 100–106, 2010.

15. Stapleton PA, McBride CR, Yi J, and Nurkiewicz TR. Uterine microvascular sensitivity to nanomaterial inhalation: An in vivo assessment. Toxicol Appl Pharmacol 288: 420–428, 2015.

16. Nurkiewicz TR, Porter DW, Barger M, Castranova V, and Boegehold MA. Particulate matter exposure impairs systemic microvascular endothelium-dependent dilation. Environ Health Perspect 112: 1299–1306, 2004.

17. Krishnan RM, Adar SD, Szpiro AA, Jorgensen NW, Van Hee VC, Barr RG, O’Neill MS, Herrington DM, Polak JF, and Kaufman JD. Vascular responses to long- and short-term exposure to fine particulate matter: MESA Air (Multi-Ethnic Study of Atherosclerosis and Air Pollution). J Am Coll Cardiol 60: 2158–2166, 2012.

18. Ragusa A, Svelato A, Santacroce C, Catalano P, Notarstefano V, Carnevali O, Papa F, Rongioletti MCA, Baiocco F, Draghi S, D’Amore E, Rinaldo D, Matta M, and Giorgini E. Plasticenta: First evidence of microplastics in human placenta. Environ Int 146: 106274, 2021.

19. Cary CM, Seymore TN, Singh D, Vayas KN, Goedken MJ, Adams S, Polunas M, Sunil VR, Laskin DL, Demokritou P, and Stapleton PA. Single inhalation exposure to polyamide micro and nanoplastic particles impairs vascular dilation without generating pulmonary inflammation in virgin female Sprague Dawley rats. Particle and Fibre Toxicology 20: 16, 2023.

20. Cary CM, Fournier SB, Adams S, Wang X, Yurkow EJ, and Stapleton PA. Single pulmonary nanopolystyrene exposure in late-stage pregnancy dysregulates maternal and fetal cardiovascular function. Toxicol Sci 199: 149–159, 2024.

21. Weiner C, Liu KZ, Thompson L, Herrig J, and Chestnut D. Effect of pregnancy on endothelium and smooth muscle: their role in reduced adrenergic sensitivity. American Journal of Physiology-Heart and Circulatory Physiology 261: H1275–H1283, 1991.

22. Magness RR, Rosenfeld CR, Hassan A, and Shaul PW. Endothelial vasodilator production by uterine and systemic arteries. I. Effects of ANG II on PGI2 and NO in pregnancy. Am J Physiol 270: H1914–1923, 1996.

23. Jansson T, and Lambert GW. Effect of intrauterine growth restriction on blood pressure, glucose tolerance and sympathetic nervous system activity in the rat at 3-4 months of age. J Hypertens 17: 1239–1248, 1999.

24. Dai Y, Zhao D, Chen CK, and Yap CH. Echocardiographic assessment of fetal cardiac function in the uterine artery ligation rat model of IUGR. Pediatr Res 90: 801–808, 2021.

25. Schreuder MF, Fodor M, van Wijk JAE, and Delemarre-van de Waal HA. Association of Birth Weight with Cardiovascular Parameters in Adult Rats During Baseline and Stressed Conditions. Pediatric Research 59: 126–130, 2006.

26. Thompson JA, Gros R, Richardson BS, Piorkowska K, and Regnault TR. Central stiffening in adulthood linked to aberrant aortic remodeling under suboptimal intrauterine conditions. Am J Physiol Regul Integr Comp Physiol 301: R1731–1737, 2011.

27. Schipke J, Gonzalez-Tendero A, Cornejo L, Willführ A, Bijnens B, Crispi F, Mühlfeld C, and Gratacós E. Experimentally induced intrauterine growth restriction in rabbits leads to differential remodelling of left versus right ventricular myocardial microstructure. Histochem Cell Biol 148: 557–567, 2017.

28. Mazzuca MQ, Tare M, Parkington HC, Dragomir NM, Parry LJ, and Wlodek ME. Uteroplacental insufficiency programmes vascular dysfunction in non-pregnant rats: compensatory adaptations in pregnancy. J Physiol 590: 3375–3388, 2012.

29. Wadley GD, McConell GK, Goodman CA, Siebel AL, Westcott KT, and Wlodek ME. Growth restriction in the rat alters expression of metabolic genes during postnatal cardiac development in a sex-specific manner. Physiol Genomics 45: 99–105, 2013.

30. Walsh SK, English FA, Johns EJ, and Kenny LC. Plasma-mediated vascular dysfunction in the reduced uterine perfusion pressure model of preeclampsia: a microvascular characterization. Hypertension 54: 345–351, 2009.

31. Stapleton PA, Nichols CE, Yi J, McBride CR, Minarchick VC, Shepherd DL, Hollander JM, and Nurkiewicz TR. Microvascular and mitochondrial dysfunction in the female F1 generation after gestational TiO2 nanoparticle exposure. Nanotoxicology 9: 941–951, 2015.

32. Orzabal MR, Lunde-Young ER, Ramirez JI, Howe SYF, Naik VD, Lee J, Heaps CL, Threadgill DW, and Ramadoss J. Chronic exposure to e-cig aerosols during early development causes vascular dysfunction and offspring growth deficits. Transl Res 207: 70–82, 2019.

33. Gandley RE, Jeyabalan A, Desai K, McGonigal S, Rohland J, and DeLoia JA. Cigarette exposure induces changes in maternal vascular function in a pregnant mouse model. Am J Physiol Regul Integr Comp Physiol 298: R1249–1256, 2010.

34. Stapleton PA, Minarchick VC, Yi J, Engels K, McBride CR, and Nurkiewicz TR. Maternal engineered nanomaterial exposure and fetal microvascular function: does the Barker hypothesis apply? Am J Obstet Gynecol 209: 227.e221-211, 2013.

35. Bird IM, Sullivan JA, D. T, Cale JM, Zhang L, Zheng J, and Magness RR. Pregnancy-Dependent Changes in Cell Signaling Underlie Changes in Differential Control of Vasodilator Production in Uterine Artery Endothelial Cells*. Endocrinology 141: 1107–1117, 2000.

36. Ni Y, Meyer M, and Osol G. Gestation increases nitric oxide–mediated vasodilation in rat uterine arteries. American Journal of Obstetrics and Gynecology 176: 856–864, 1997.

37. Nelson SH, Steinsland OS, Wang Y, Yallampalli C, Dong YL, and Sanchez JM. Increased nitric oxide synthase activity and expression in the human uterine artery during pregnancy. Circ Res 87: 406–411, 2000.

38. Boujedaini N, Liu J, Thuillez C, Cazin L, and Mensah-Nyagan AG. In vivo regulation of vasomotricity by nitric oxide and prostanoids during gestation. Eur J Pharmacol 427: 143–149, 2001.

39. Bird IM, Zheng J, Cale JM, and Magness RR. Pregnancy induces an increase in angiotensin II type-1 receptor expression in uterine but not systemic artery endothelium. Endocrinology 138: 490–498, 1997.

40. Rennie MY, Rahman A, Whiteley KJ, Sled JG, and Adamson SL. Site-Specific Increases in Utero- and Fetoplacental Arterial Vascular Resistance in eNOS-Deficient Mice Due to Impaired Arterial Enlargement1. Biology of Reproduction 92: 2015.

41. Chang JK, Roman C, and Heymann MA. Effect of endothelium-derived relaxing factor inhibition on the umbilical-placental circulation in fetal lambs in utero. Am J Obstet Gynecol 166: 727–734, 1992.

42. Kusinski LC, Stanley JL, Dilworth MR, Hirt CJ, Andersson IJ, Renshall LJ, Baker BC, Baker PN, Sibley CP, Wareing M, and Glazier JD. eNOS knockout mouse as a model of fetal growth restriction with an impaired uterine artery function and placental transport phenotype. Am J Physiol Regul Integr Comp Physiol 303: R86–93, 2012.

43. Neerhof MG, Synowiec S, Khan S, and Thaete LG. Impact of endothelin A receptor antagonist selectivity in chronic nitric oxide synthase inhibition-induced fetal growth restriction in the rat. Hypertens Pregnancy 29: 284–293, 2010.

44. Thaete LG, Kushner DM, Dewey ER, and Neerhof MG. Endothelin and the regulation of uteroplacental perfusion in nitric oxide synthase inhibition-induced fetal growth restriction. Placenta 26: 242–250, 2005.

45. Gokina NI, Mandalà M, and Osol G. Induction of localized differences in rat uterine radial artery behavior and structure during gestation. Am J Obstet Gynecol 189: 1489–1493, 2003.

46. Rennie MY, Whiteley KJ, Adamson SL, and Sled JG. Quantification of Gestational Changes in the Uteroplacental Vascular Tree Reveals Vessel Specific Hemodynamic Roles During Pregnancy in Mice. Biol Reprod 95: 43, 2016.

47. Nurkiewicz TR, Porter DW, Hubbs AF, Stone S, Chen BT, Frazer DG, Boegehold MA, and Castranova V. Pulmonary nanoparticle exposure disrupts systemic microvascular nitric oxide signaling. Toxicol Sci 110: 191–203, 2009.

48. LeBlanc AJ, Moseley AM, Chen BT, Frazer D, Castranova V, and Nurkiewicz TR. Nanoparticle inhalation impairs coronary microvascular reactivity via a local reactive oxygen species-dependent mechanism. Cardiovasc Toxicol 10: 27–36, 2010.

49. Wauters A, Dreyfuss C, Pochet S, Hendrick P, Berkenboom G, van de Borne P, and Argacha JF. Acute exposure to diesel exhaust impairs nitric oxide-mediated endothelial vasomotor function by increasing endothelial oxidative stress. Hypertension 62: 352–358, 2013.

50. Cherng TW, Paffett ML, Jackson-Weaver O, Campen MJ, Walker BR, and Kanagy NL. Mechanisms of diesel-induced endothelial nitric oxide synthase dysfunction in coronary arterioles. Environ Health Perspect 119: 98–103, 2011.

51. El-Mahdy MA, Abdelghany TM, Hemann C, Ewees MG, Mahgoup EM, Eid MS, Shalaan MT, Alzarie YA, and Zweier JL. Chronic cigarette smoke exposure triggers a vicious cycle of leukocyte and endothelial-mediated oxidant stress that results in vascular dysfunction. Am J Physiol Heart Circ Physiol 319: H51–h65, 2020.

52. Minarchick VC, Stapleton PA, Porter DW, Wolfarth MG, çiftyürek E, Barger M, Sabolsky EM, and Nurkiewicz TR. Pulmonary cerium dioxide nanoparticle exposure differentially impairs coronary and mesenteric arteriolar reactivity. Cardiovasc Toxicol 13: 323–337, 2013.

53. Cary CM, Seymore TN, Singh D, Vayas KN, Goedken MJ, Adams S, Polunas M, Sunil VR, Laskin DL, Demokritou P, and Stapleton PA. Single inhalation exposure to polyamide micro and nanoplastic particles impairs vascular dilation without generating pulmonary inflammation in virgin female Sprague Dawley rats. Part Fibre Toxicol 20: 16, 2023.

54. Stapleton PA, McBride CR, Yi J, Abukabda AB, and Nurkiewicz TR. Estrous cycle-dependent modulation of in vivo microvascular dysfunction after nanomaterial inhalation. Reprod Toxicol 78: 20–28, 2018.

55. Moreno GM, Brunson-Malone T, Adams S, Nguyen C, Seymore TN, Cary CM, Polunas M, Goedken MJ, and Stapleton PA. Identification of micro- and nanoplastic particles in postnatal sprague-dawley rat offspring after maternal inhalation exposure throughout gestation. Sci Total Environ 951: 175350, 2024.

56. Fournier SB, Kallontzi S, Fabris L, Love C, and Stapleton PA. Effect of Gestational Age on Maternofetal Vascular Function Following Single Maternal Engineered Nanoparticle Exposure. Cardiovasc Toxicol 19: 321–333, 2019.

57. Bendall JK, Douglas G, McNeill E, Channon KM, and Crabtree MJ. Tetrahydrobiopterin in cardiovascular health and disease. Antioxid Redox Signal 20: 3040–3077, 2014.

58. Krejsa CM, Franklin CC, White CC, Ledbetter JA, Schieven GL, and Kavanagh TJ. Rapid activation of glutamate cysteine ligase following oxidative stress. J Biol Chem 285: 16116–16124, 2010.

59. Gilbert JS, Bauer AJ, Gingery A, Banek CT, and Chasson S. Circulating and utero-placental adaptations to chronic placental ischemia in the rat. Placenta 33: 100–105, 2012.

60. McClements L, Richards C, Patel N, Chen H, Sesperez K, Bubb KJ, Karlstaedt A, and Aksentijevic D. Impact of reduced uterine perfusion pressure model of preeclampsia on metabolism of placenta, maternal and fetal hearts. Sci Rep 12: 1111, 2022.

61. Payne JA, Alexander BT, and Khalil RA. Reduced endothelial vascular relaxation in growth-restricted offspring of pregnant rats with reduced uterine perfusion. Hypertension 42: 768–774, 2003.

62. Barron C, Mandala M, and Osol G. Effects of pregnancy, hypertension and nitric oxide inhibition on rat uterine artery myogenic reactivity. J Vasc Res 47: 463–471, 2010.

63. Roggensack AM, Zhang Y, and Davidge ST. Evidence for Peroxynitrite Formation in the Vasculature of Women With Preeclampsia. Hypertension 33: 83–89, 1999.

64. Lauer T, Preik M, Rassaf T, Strauer BE, Deussen A, Feelisch M, and Kelm M. Plasma nitrite rather than nitrate reflects regional endothelial nitric oxide synthase activity but lacks intrinsic vasodilator action. Proc Natl Acad Sci U S A 98: 12814–12819, 2001.

65. Herold S, Exner M, and Boccini F. The Mechanism of the Peroxynitrite-Mediated Oxidation of Myoglobin in the Absence and Presence of Carbon Dioxide. Chemical Research in Toxicology 16: 390–402, 2003.

66. Nurkiewicz TR, Porter DW, Hubbs AF, Stone S, Moseley AM, Cumpston JL, Goodwill AG, Frisbee SJ, Perrotta PL, Brock RW, Frisbee JC, Boegehold MA, Frazer DG, Chen BT, and Castranova V. Pulmonary particulate matter and systemic microvascular dysfunction. Res Rep Health Eff Inst 3-48, 2011.

67. Du Y, Navab M, Shen M, Hill J, Pakbin P, Sioutas C, Hsiai TK, and Li R. Ambient ultrafine particles reduce endothelial nitric oxide production via S-glutathionylation of eNOS. Biochemical and Biophysical Research Communications 436: 462–466, 2013.

68. Förstermann U, and Münzel T. Endothelial Nitric Oxide Synthase in Vascular Disease. Circulation 113: 1708–1714, 2006.

69. Janaszak-Jasiecka A, Płoska A, Wierońska JM, Dobrucki LW, and Kalinowski L. Endothelial dysfunction due to eNOS uncoupling: molecular mechanisms as potential therapeutic targets. Cell Mol Biol Lett 28: 21, 2023.

70. Kukreja RC, and Xi L. eNOS phosphorylation: a pivotal molecular switch in vasodilation and cardioprotection? J Mol Cell Cardiol 42: 280–282, 2007.

71. McCabe TJ, Fulton D, Roman LJ, and Sessa WC. Enhanced electron flux and reduced calmodulin dissociation may explain “calcium-independent” eNOS activation by phosphorylation. J Biol Chem 275: 6123–6128, 2000.

72. Ying Z, Xu X, Chen M, Liu D, Zhong M, Chen LC, Sun Q, and Rajagopalan S. A synergistic vascular effect of airborne particulate matter and nickel in a mouse model. Toxicol Sci 135: 72–80, 2013.

73. Abdelghany TM, Ismail RS, Mansoor FA, Zweier JR, Lowe F, and Zweier JL. Cigarette smoke constituents cause endothelial nitric oxide synthase dysfunction and uncoupling due to depletion of tetrahydrobiopterin with degradation of GTP cyclohydrolase. Nitric Oxide 76: 113–121, 2018.

74. Ying Z, Kampfrath T, Thurston G, Farrar B, Lippmann M, Wang A, Sun Q, Chen LC, and Rajagopalan S. Ambient Particulates Alter Vascular Function through Induction of Reactive Oxygen and Nitrogen Species. Toxicological Sciences 111: 80–88, 2009.

75. Chen C-A, Wang T-Y, Varadharaj S, Reyes LA, Hemann C, Talukder MAH, Chen Y-R, Druhan LJ, and Zweier JL. S-glutathionylation uncouples eNOS and regulates its cellular and vascular function. Nature 468: 1115–1118, 2010.

76. Landmesser U, Dikalov S, Price SR, McCann L, Fukai T, Holland SM, Mitch WE, and Harrison DG. Oxidation of tetrahydrobiopterin leads to uncoupling of endothelial cell nitric oxide synthase in hypertension. J Clin Invest 111: 1201–1209, 2003.

77. Hayashi T, Fukuto JM, Ignarro LJ, and Chaudhuri G. Basal release of nitric oxide from aortic rings is greater in female rabbits than in male rabbits: implications for atherosclerosis. Proc Natl Acad Sci U S A 89: 11259–11263, 1992.

78. Osol G, Celia G, Gokina N, Barron C, Chien E, Mandala M, Luksha L, and Kublickiene K. Placental growth factor is a potent vasodilator of rat and human resistance arteries. Am J Physiol Heart Circ Physiol 294: H1381–1387, 2008.

79. Tropea T, De Francesco EM, Rigiracciolo D, Maggiolini M, Wareing M, Osol G, and Mandalà M. Pregnancy Augments G Protein Estrogen Receptor (GPER) Induced Vasodilation in Rat Uterine Arteries via the Nitric Oxide - cGMP Signaling Pathway. PLoS One 10: e0141997, 2015.

80. Storment JM, Meyer M, and Osol G. Estrogen augments the vasodilatory effects of vascular endothelial growth factor in the uterine circulation of the rat. Am J Obstet Gynecol 183: 449–453, 2000.

81. Burger NZ, Kuzina OY, Osol G, and Gokina NI. Estrogen replacement enhances EDHF-mediated vasodilation of mesenteric and uterine resistance arteries: role of endothelial cell Ca2+. Am J Physiol Endocrinol Metab 296: E503–512, 2009.

82. Fournier SB, D’Errico JN, Adler DS, Kollontzi S, Goedken MJ, Fabris L, Yurkow EJ, and Stapleton PA. Nanopolystyrene translocation and fetal deposition after acute lung exposure during late-stage pregnancy. Part Fibre Toxicol 17: 55, 2020.

