## Supplemental Table 1 for "Micro and nanoplastic inhalation during pregnancy elicits uterine endothelial dysfunction in Sprague Dawley rats by impeding nitric oxide signaling"

| Target | Manufacturer | Catalog Numbe | dilution | Host species |
|---|---|---|---|---|
| **GTPCH1** | Invitrogen | pa5-103865 | 1:1000 | rabbit |
| **eNOS** | Cell Signaling Technology | 32027s | 1:500 | rabbit |
| **phospho-eNOS (Ser1177)** | Invitrogen | MA5-14957 | 1:1000 | rabbit |
| **Arginase 1** | Proteintech | 161001-1-AP | 1:1000 | rabbit |
| **iNOS** | Proteintech | 18985-1-AP | 1:1000 | rabbit |
| **Thioredoxin** | Cell Signaling Technology | 2429S | 1:1000 | rabbit |
| **3-Nitrotyrosine** | Abcam | AB61392 | 1:1000 | mouse |
| **DHFR** | Abcam | ab124814 | 1:1000 | rabbit |
| **GCLC (glutamate cysteine ligas** | Abcam | ab190685 | 1:1000 | rabbit |
| **Beta-actin** | Cell Signaling Technology | 3700S | 1:1000 | mouse |

Table S1. Antibody sources and concentrations utilized for Western Blot analyses.
