## Supplementary figures and images for "Micro and nanoplastic inhalation during pregnancy elicits uterine endothelial dysfunction in Sprague Dawley rats by impeding nitric oxide signaling"

### Supplemental Figure 1

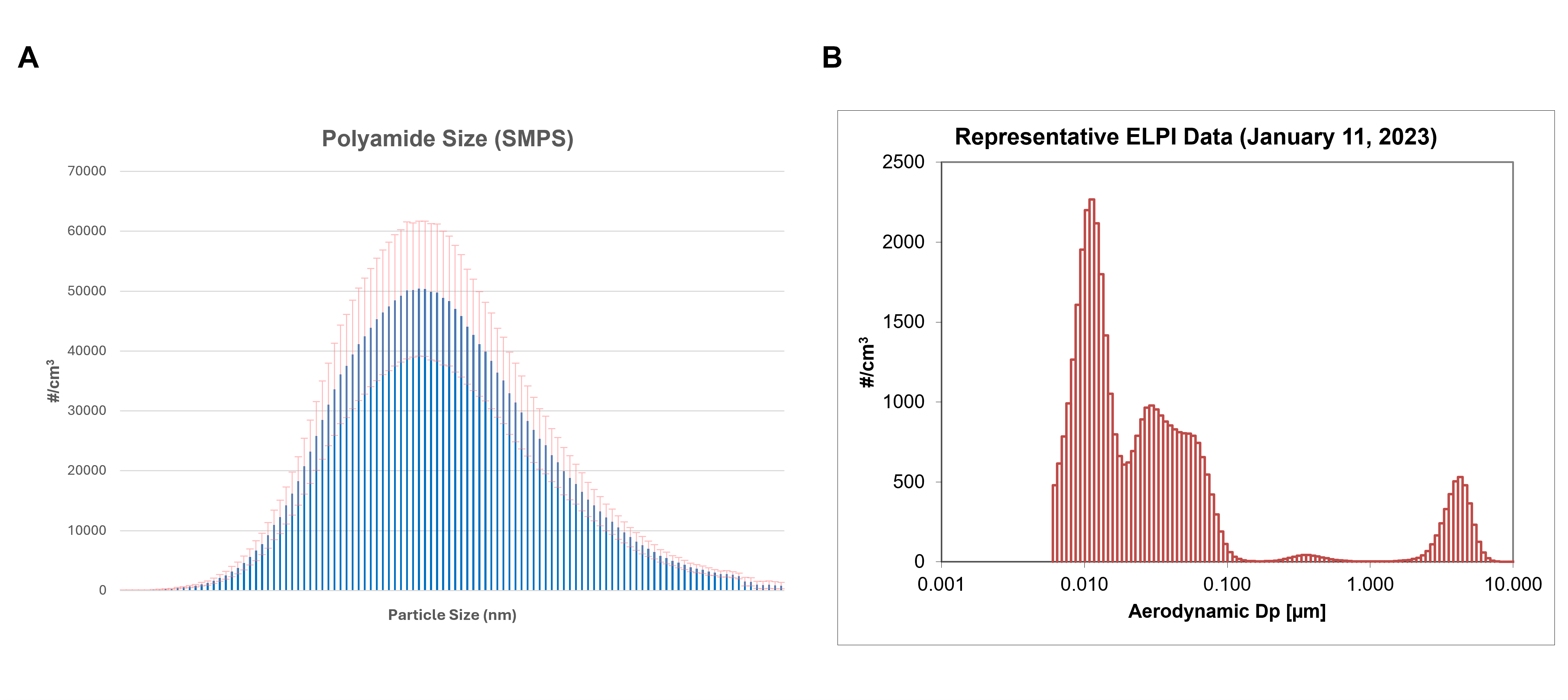

### Supplemental Figure 2

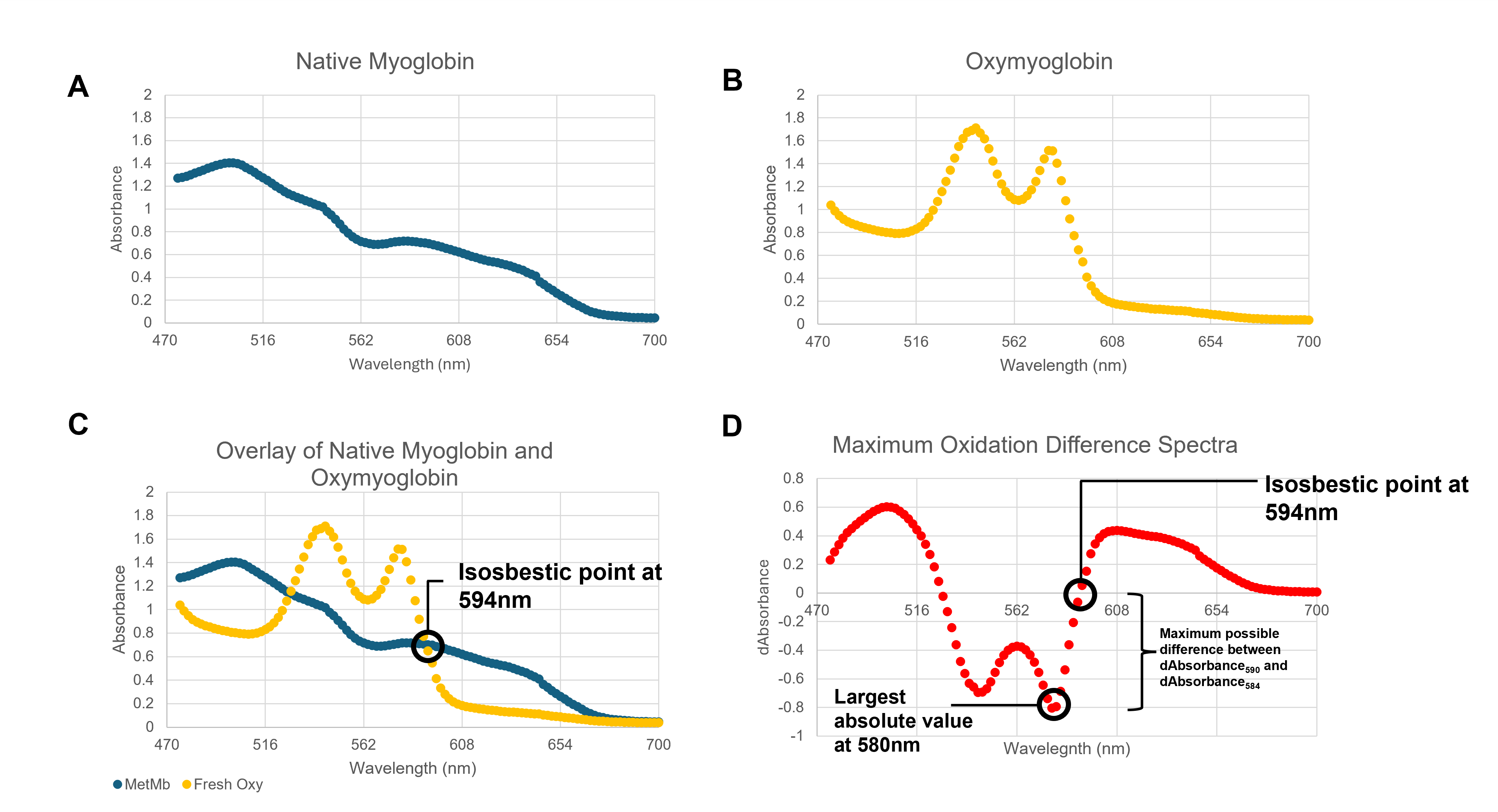
