## Supplemental Table 2 for "Micro and nanoplastic inhalation during pregnancy elicits uterine endothelial dysfunction in Sprague Dawley rats by impeding nitric oxide signaling"

| Group | N | n | Active Internal Diameter (µm | Maximum Internal Diameter (µm) | Vessel Tone (%) |
|---|---|---|---|---|---|
| Control | 20 | 22 | 118± 10.1 | 194 ± 10.9 | 40.5 ± 3.15 |
| Polyamide | 28 | 30 | 112 ± 6.5 | 185 ± 7.7 | 37.8 ± 2.46 |

Table S2. Vessel characteristics of radial arteries. Values are presented as mean ± SEM. “N” is representative of the number of animals, “n” is representative of the number of vessels. Significant differences ($p < 0.05$) from the control group are denoted by *.
